# Elevated plasma pTau181 in a specialty neuropsychiatric clinic is associated with opioid use and guides medication titration on a clinically relevant short-time scale

**DOI:** 10.1101/2024.12.01.626163

**Authors:** Emily Hanson, Karl Berner, Jon Berner

## Abstract

A combinatorial explosion of agents that delay neurodegeneration in animal models has created a crisis of abundance in human applications with too many combinations to try when time is short. Plasma biomarkers of premature neuronal aging, the amyloid-tau-neurodegeneration (ATN) profile, may provide a tool to rapidly optimize treatment in single-patient trials. We retrospectively analyzed plasma ATN profiles in patients with extensive neuropsychiatric polypharmacy and premature neuronal aging. ATN profiles were based on the biomarkers amyloid-β ratio, phosphorylated Tau 181 (pTau181), and neurofilament light chain (NfL). We investigated whether ATN profile biomarkers were associated with age, gender, metabolic syndrome markers, and medication use. Two case reports additionally provide examples of the application of ATN profiles in clinical settings. We found that in 80 patients with ATN profiles and complete clinical phenotypes, pTau181 was elevated in 31 (38.75%), amyloid-β ratio was below normal ranges in 11 (13.75%), and NfL was elevated in three (3.75%). The biomarkers correlated with age, as expected. Opioid use was significantly associated with pTau181 (p=0.004) and NfL (p=0.002), also after Bonferroni correction (both p <0.05), but not with amyloid-β ratio. No other medications were associated with the biomarkers. In conclusion, identifying the overlaps between complex behavioral phenotypes (pain and cognition), plasma endophenotypes (ATN profile), and medication-targeted components of age-related pathophysiology is now a technical problem rather than theoretical speculation. Rapid progress in single-patient trials for treatment optimization will require funding to promote using repurposed generic treatments, educate patients and providers regarding optimization principles, and continue developing sensitive biomarkers.

## Introduction

Combinatorial treatments with multiple medications are uniformly most effective in slowing age-related neurodegeneration in short-lived animal models of disease, consistent with the diversity of pathological aging pathways [1]. Recent data in the model organism *Drosophila melanogaster* show additive and thus orthogonal substantial effects of lithium and rapamycin in combination (30% lifespan extension) [2]. This finding is particularly interesting because these drugs are inexpensive and managed with a reasonable safety profile in diverse human populations.

All ongoing human trials of rapamycin and lithium are limited to evaluating single agents, typically over long treatment times. As a result, treatment with a combination of rapamycin and lithium to promote health during aging has not been tested in clinical trials yet [3]. There are various barriers to researching such treatments, including the generic status of these medications without industry financial support, the minimal availability of a clinical labor force with intersectional training, and perceived regulatory monitoring compliance costs. Until recently, an additional constraint in this field was the difficulty in gathering tissue data in a cost-effective manner with a time constant of response allowing for multiple medication adjustments within the lifespan of a patient. That is, analyzing cerebrospinal fluid (CSF) is invasive and expensive, detecting changes in magnetic resonance imaging volume is slow and has small effect sizes, and performing positron emission tomography scans is expensive while access is geographically limited.

The recent commercial introduction of tests for plasma biomarkers of neurodegeneration allows for more rapid optimization of prophylactic treatments against neurodegeneration. The amyloid-tau-neurodegeneration (ATN) profile is determined in plasma based on the biomarkers amyloid-β ratio, phosphorylated Tau 181 (pTau181), and neurofilament light chain (NfL) and is used to evaluate age-related mild cognitive decline. The presence of amyloid-β plaques and intracellular Tau aggregates in the brain are hallmarks of several neurodegenerative diseases, and plasma NfL is a non-specific marker of neuronal injury [4, 5]. Extensive research has shown the relevance of the ATN profile for the diagnosis of Alzheimer’s Disease (AD) and AD-related dementias [5]. The ATN profile may allow for multiple treatment trials in an individual patient over a span of one to five years, a time span more congruent with clinical needs associated with neurodegeneration prophylaxis.

To determine the value of ATN profiles in guiding management in our psychiatric clinic, we performed a retrospective analysis of the ATN profile in a cohort of patients with premature neuronal aging. We then assessed whether medication use was associated with any of the biomarkers of the ATN profile. In addition, we described the implications of the ATN profile in two case reports of patients with premature neuronal aging in treatment with lithium or rapamycin.

## Methods

### Study population

Patients who spontaneously presented with subjective cognitive complaints and for whom ATN profiles were obtained as part of their diagnostic assessment were recruited from a psychiatric practice. Patients were included in this retrospective study between February 17^th^, 2024, and July 3^rd^, 2024.

Clinical data were collected from chart review such as age, gender, height, weight, body mass index (BMI), fasting blood sugar (FBS), triglyceride to high-density lipoprotein (TG/HDL) ratio, insulin to BMI ratio, red cell distribution width (RDW), neutrophil to lymphocyte ratio, whether the patient had tried more than one anti-depressant, and which psychiatric medications (i.e., lithium, benzodiazepine, antipsychotic, anticonvulsant, opioid, ketamine, psychostimulant) the patient was on during their ATN testing.

### Informed consent

This retrospective analysis of existing data derived from standard clinical practice was exempt from Institutional Review Board approval under Category 2 of the Basic Health and Human Services Policy for Protection of Human Research Subjects Subpart A Section 46.101 [6]. No informed consent was required.

### Sample collection

Non-fasting blood samples were collected through venipuncture into Ethylenediaminetetraacetic Acid (EDTA) tubes. Plasma was separated from blood within 40 minutes after collection by centrifugation at >1500g for 10 minutes. It was then transferred to two LabCorp polypropylene transport tubes (LabCorp, Burlington, NC) and sealed. Plasma samples were stored up to 24 hours at -20°C before shipment on dry ice to LabCorp.

### ATN profile measurements

Concentrations of amyloid-β 42 (Aβ42) and amyloid-β40 (Aβ40) were measured with a chemiluminescence enzyme immunoassay (CLEIA) based on Sysmex reagents and technology (Sysmex, Kobe, Japan). The amyloid-β ratio (Aβ42/Aβ40) was subsequently calculated. A normal amyloid-β ratio is >0.102. Concentrations of pTau181 and NfL were analyzed using electrochemiluminescence immunoassays (ECLIAs) based on Roche reagents and technology (Roche Diagnostics, Indianapolis, IN). Normal pTau181 range for individuals 20–55 years of age: 0.00–0.95 pg/ml, normal range for individuals >55 years of age: 0.00–0.97 pg/ml. Normal NfL range: 0.00–2.13 pg/ml.

### Statistical analysis

Patient data were entered into an Excel database (Microsoft, Redmond, WA) and exported into GNU PSPP (https://www.gnu.org/software/pspp/) for data analysis. Variables are presented as mean with standard deviation (SD), median, range, or proportion. Pearson correlation coefficients (r) were calculated between ATN variables and clinical phenotypes. Bonferroni adjustment was performed post-hoc (for 8 comparisons) on two-tailed statistical findings related to current medication use. A two-tailed p-value <0.05 was considered significant.

## Results

### Patients

In total, 113 patients with premature neuronal aging ATN profiles were identified through the chart reviews. The mean age of these patients was 60.4 (SD 12.3) years, and the median age was 61 years. The majority of these patients were female (n = 77/113, 68%). The patients had a mean BMI of 28.4 (SD 6.3) kg/m^2^. Clinical data are presented in **Table 1**. Of these patients, 36 had tried >1 anti-depressant. At the time of the blood draw for ATN testing, the medications they were using were lithium (n=12), benzodiazepine (n=61), antipsychotic (n =36), anticonvulsant (n=53), opioid (n=18), ketamine (n=22), psychostimulant (n=45).

**Table 1.** Clinical data of N = 113 patients.

|  | **Mean (SD)** | **Median** | **Minimum** | **Maximum** | **Range** |
| --- | --- | --- | --- | --- | --- |
| **Age, in years** | 60.4 (12.3) | 61 | 23 | 92 | 69 |
| **Height<sup>\*</sup>, in cm** | 168.9 (10.5) | 167.6 | 142.6 | 198.1 | 55.9 |
| **Weight<sup>\*\*</sup>, in kg** | 81.1 (20.3) | 79.8 | 44.9 | 162.8 | 117.9 |
| **BMI, in kg/m<sup>2</sup>** | 28.4 (6.3) | 27.2 | 17.6 | 50.1 | 32.4 |
| **FBS, in mg/dL** | 97.9 (19.7) | 95 | 67 | 164 | 97 |
| **TG/HDL ratio** | 2.8 (2.3) | 1.9 | 0.5 | 11 | 10.4 |
| **insulin to BMI ratio** | 2.1 (1.4) | 1.5 | 0.1 | 6 | 6 |
| **RDW, in %** | 13.2 (1.7) | 12.9 | 11.7 | 21.1 | 9.4 |
| **neutrophil to lymphocyte ratio** | 2.1 (0.9) | 1.9 | 0.8 | 4.4 | 3.5 |

\*Height missing for 14 patients, \*\*Weight missing for 10 patients.
BMI, body mass index; FBS, fasting blood sugar; RDW, red cell distribution width; SD, standard deviation; TG/HDL, triglyceride to high-density lipoprotein ratio.

### ATN profile biomarkers were correlated with age

Of 80 of the 113 patients with a full phenotypical profile, the ATN profile had been determined in plasma. The mean pTau181 concentration was 0.93 (SD 0.35) pg/ml, the mean NfL concentration was 2.85 (SD 1.97) pg/ml, and the mean amyloid-β ratio was 0.1 (SD 0.01) (**Table 2**). Of these patients, 31 (38.75%) had a pTau181 concentration above normal ranges, 11 (13.75%) had an amyloid-β ratio below normal ranges, and three (3.75%) had an NfL concentration above normal ranges. None of the 80 patients tested had all three values in abnormal ranges.

**Table 2.** ATN profiles in n = 80 patients.

|  | **Mean (SD)** | **Range** | **Patients, n (%)** |
| --- | --- | --- | --- |
| **pTau181, in pg/ml** | 0.93 (0.35) | 0.48–2.26 |  |
| pTau181 within normal ranges | 0.71 (0.13) | 0.45–0.93 | 51 (63.75%) |
| pTau181 above normal ranges | 1.3 (0.28) | 0.98–2.26 | 29 (36.25%) |
| **Aβ42, in pg/ml** | 19.9 (4.4) | 9.7–35.8 |  |
| **Aβ40, in pg/ml** | 175.5 (33.2) | 91.5–188.9 |  |
| **amyloid-β ratio** | 0.1 (0.01) | 0.09–0.14 |  |
| amyloid-β ratio within normal ranges | 0.12 (0.01) | 0.10–0.14 | 69 (86.25%) |
| amyloid-β ratio below normal ranges | 0.096 (0.005) | 0.089–0.10 | 11 (13.75%) |
| **NfL, in pg/ml** | 2.85 (1.97) | 0.56–12.9 |  |
| NfL within normal ranges | 2.54 (1.14) | 0.56–6.9 | 77 (96.25%) |
| NfL above normal ranges | 10.79 (2.39) | 8.19–12.9 | 3 (3.75%) |

Aβ40, $\beta$ -amyloid 40; Aβ 42, $\beta$ -amyloid 42; ATN, amyloid-Tau-neurodegeneration; NfL, neurofilament light chain; n, number; pTau181, phosphorylated Tau 181; SD, standard deviation.

Age was found to be correlated with pTau181 concentration (r = 0.405, p <0.000), NfL concentration (r = 0.384, p <0.000), and amyloid-β ratio (r = -0.285, p <0.010) (**Figure 1**).

**Figure 1.**
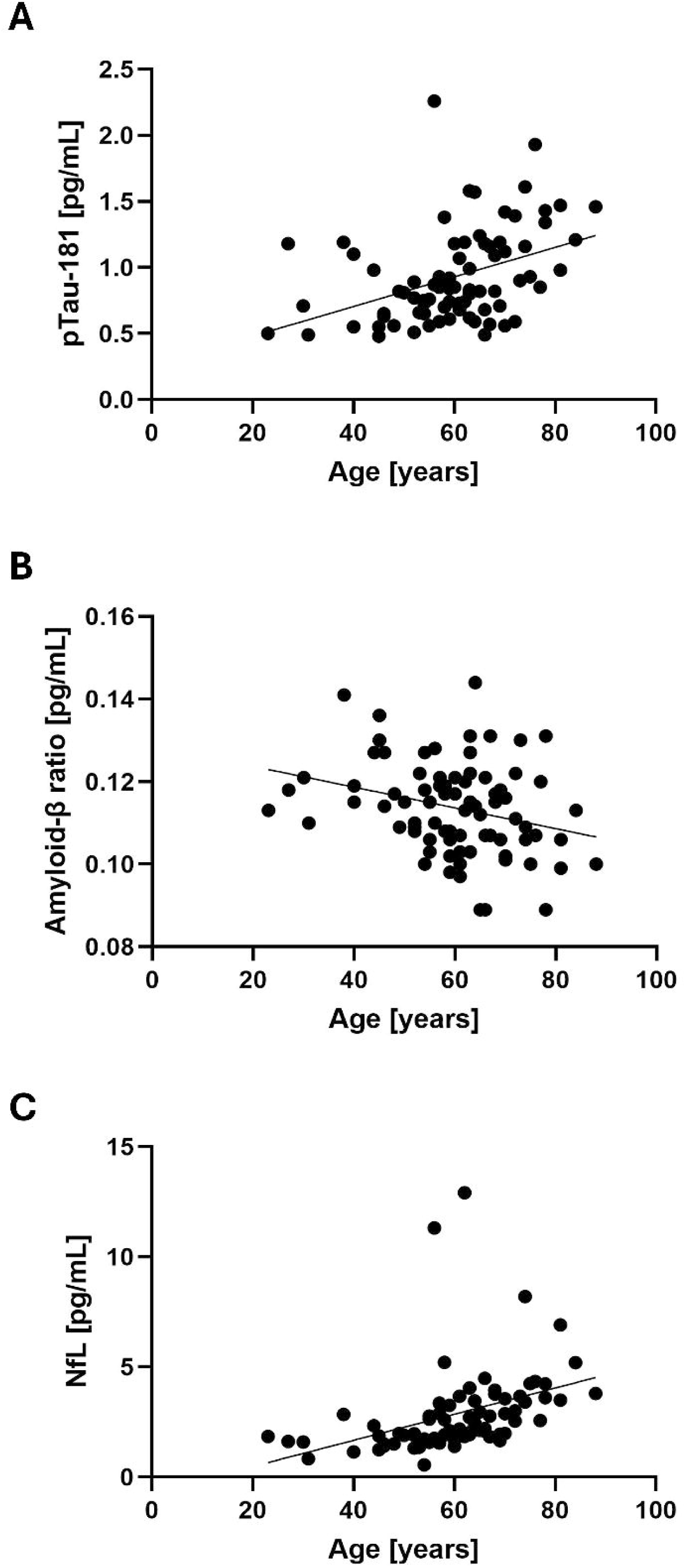
**Correlations between ATN profile biomarkers and age** Scatter plots of the ATN profile biomarkers and age of the patients (n=80). Each black dot represents a single value in one patient, the thin line represents the simple linear regression. **A** Correlation between pTau181 concentration and age (r = 0.405, p <0.000). **B** Correlation between NfL concentration and age (r = 0.384, p <0.000). **C** Correlation between amyloid-β ratio and age (r = -0.285, p <0.010). ATN, amyloid-tau-neurodegeneration; NfL, neurofilament light chain; pTau181, phosphorylated Tau 181.

### ATN profile biomarkers were not correlated with metabolic syndrome markers

As the metabolic syndrome markers FBS, TG/HDL ratio, and insulin to BMI ratio were not available for all 80 patients, potential association with biomarkers of the ATN profile was analyzed in smaller subsets. ATN profile biomarkers were not associated with any of the metabolic syndrome markers in the subset of patients with a full metabolic profile (see **Supplemental Figure S1**).

### Opioid use was associated with pTau181 and NfL

We assessed associations between current medication use or previously failed medication(s) and the biomarkers of the ATN profile. We found that opioid use was significantly associated with pTau181 concentration (r = 0.315, p = 0.004) (**Figure 2A**) and NfL concentration (r = 0.341, p = 0.002) (**Figure 2B**), but not with the amyloid-β ratio (r = 0.157, p = 0.165) (**Figure 2C**). After Bonferroni correction for multiple comparisons across eight medication classes, pTau181 and NfL concentrations were still significantly associated with opioid use (p <0.05).

**Figure 2.**
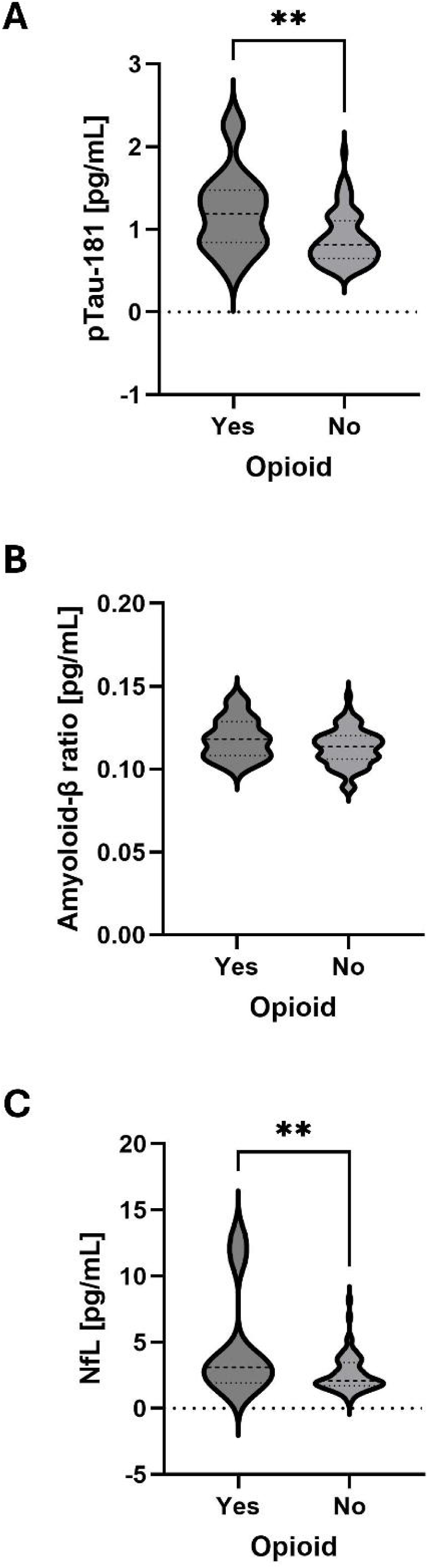
**Opioid use was associated with higher pTau181 and NfL** Violin plots of ATN profile biomarkers in patients (n=80). The dashed line in the violin plot indicates the median, the dotted lines indicate the quartiles. **A** pTau181 concentration. **B** NfL concentration. **C** amyloid-β ratio. ** p< 0.005. ATN, amyloid-tau-neurodegeneration; NfL, neurofilament light chain; pTau181, phosphorylated Tau 181.

Psychostimulant use was significantly, but weakly, associated with the amyloid-β ratio (r = 0.023, p = 0.039). After Bonferroni correction, this finding was no longer significant (p >0.05). The other medications and previous medication failure(s) were not associated with any of the biomarkers of the ATN profile.

### Case report #1, lithium treatment

A 65-year-old female patient presented in March 2021, at 61 years old, for bipolar disorder. Her disorder remained stable on lamotrigine 100 mg and Adderall 30 mg until February 2024. At that time, given the progression of irritability and concentration disturbance, in combination with dense penetrance of Parkinson’s disease in her mother’s line, her baseline ATN profile was assessed. Her pTau181 and NfL concentrations were above the normal range, and her amyloid-β ratio was below the normal range. Based on her ATN profile, the patient elected to start with lithium. Subsequently, over the next six months, her irritability resolved (reported by patient and spouse) and her pTau181 concentrations decreased to within the normal range for her age. Longitudinal data of her pTau181 concentrations are presented in **Table 3** and **Supplemental Figure S2**.

**Table 3.** ATN profile of case report 1.

| **day** | **Lithium dose, in mg** | **amyloid-β ratio** | **Aβ42, in pg/ml** | **Aβ40, in pg/ml** | **pTau181, in pg/ml** | **NfL, in pg/ml** |
| --- | --- | --- | --- | --- | --- | --- |
| **0** | 0 | 0.089 | 14.28 | 160.73 | 1.24 | 2.96 |
| **97** | 300* | 0.099 | 17.24 | 173.79 | 1.18 | 3.22 |
| **193** | 450** | 0.094 | 12.74 | 135.11 | 0.95 | 2.98 |

Aβ40, $\beta$ -amyloid 40; Aβ 42, $\beta$ -amyloid 42; ATN, amyloid-Tau-neurodegeneration; NfL, neurofilament light chain; n, number; pTau181, phosphorylated Tau 181.
\*since day 23. \*\*since day 107.

### Case report #2, rapamycin prophylaxis

A 63-year-old female patient has been seen in our clinic since 2006. Initially, she presented with a non-specific mood disorder that was treated with 225 mg venlafaxine, and leukoencephalopathy with an associated pain syndrome that was treated with transdermal fentanyl 25 μg/hr. Her subsequent course has been insidiously progressive over 18 years with the emergence of atonic seizures, bipolar disorder, Parkinson’s disease, and relentless progression of neurogenic pain. Her mother also has Parkinson’s disease. Multiple sclerosis was ruled out by numerous evaluations. Multivariate analysis of her CSF metabolome revealed substantial z-decrements in homocarnosine suggestive of accelerated astroglial aging/inflammation (see patient number #895351 in the first figure of [7]).

Rapamycin 6 mg weekly was initiated in June 2020 and subsequently titrated to effect over multiple years to 4 mg. During the titration period, she considered suicide due to poor pain control despite 80 mg oxycodone. Her dominant response to rapamycin was near complete resolution of fatigue which formerly kept her bedbound most of her day. Her secondary response, likely in conjunction with acarbose 300 mg daily (prescribed as prophylaxis for opioid constipation and dementia) and rotation to 16 mg of buprenorphine was “perfect” pain control. Despite treatment, her pTau181 concentration was 1.07 pg/ml in March 2024, with subsequent longitudinal monitoring planned. Despite superficial disease remission, informed consent for further augmentation trials is required given ongoing objective active disease based on the ATN profile biomarkers. Similar surveillance protocols for patients at high risk of relapse are standard of care in diverse clinical settings, e.g., elevated serum cancer biomarkers in Oncology and serum inflammatory biomarkers in Rheumatology.

## Discussion

The chief finding in this retrospective study was that opioid use was significantly associated with elevated concentrations of the neurodegeneration markers pTau181 and NfL in plasma. An additional incidental finding was that high pTau181 had immediate relevance for the clinical management of the anti-aging compounds lithium and rapamycin.

The association between opioid use and high plasma pTau181 was consistent with a large study (n=995) that discovered an association between chronic pain and increased pTau181 in CSF. That study also found an association between chronic pain, total Tau, and a CSF marker (TNF) of “M1-like” microglial activation [8]. Note that the community study, which included subjects with mild cognitive impairment, subjects with AD, as well as healthy controls, likely sampled an overall less sick population, and did not determine axonal degeneration associated with pain as they did not analyze NfL concentration [8]. The preliminary case reports we presented here suggest that lithium and rapamycin may, in select individuals, treat subjective pain complaints and objective markers of central nervous system inflammation.

Chronic lithium use for dementia prophylaxis is massively underutilized in community settings despite double-blind studies documenting efficacy in humans [9], and epidemiological studies linking variable dementia prevalences to trace lithium concentrations in the water supply [10]. Optimal lithium titration for dementia prophylaxis is limited by a host of subtle side effects well-known to experienced psychiatric providers [11], strongly arguing against guidelines based on population averages rather than single-patient trials using ATN profiles to guide optimal titration.

Trials of rapamycin for dementia prophylaxis are ongoing [12], and one small human trial (n=115) showed effective prophylaxis against age-related pain in females after 48 weeks on a very low dose of rapamycin [13]. Optimal dose titration of rapamycin in the individual patient is even less well-studied than dose titration of lithium, with almost absent structured clinical data regarding side effects between doses typically used for healthy longevity patients (i.e., 1 mg) [14] and the higher doses (i.e., 3-5 mg) used for transplant rejection prophylaxis or systemic lupus erythematosus [15].

The major limitation of this retrospective study design is the sampling bias of the estimated effect. The small effect size of the associations between the two biomarkers pTau181 and NfL and opioid use, given that r squared is approximately 10% of the variance, questions the clinical utility of this plasma biomarker to guide management in isolation. Furthermore, causal inferences regarding the risk vs. benefit of chronic opioid use cannot be derived from a cross-sectional design with non-randomized sampling. The “true” population response to chronic opioid use is very likely highly constrained by psychosocial factors. In Washington state, chronic opioid use is highly stigmatized and regulated with minimal patient access unless a patient has substantial financial resources and geographic proximity to available providers. In addition, providers are hesitant to prescribe opioids due to poorly defined patient characteristics in combination with working in a de facto strict liability regime in case of an overdose death of any type. Only randomized trials or longitudinal single-patient trials will allow for causal inferencing based on clinically relevant effect sizes, a challenging task in today’s regulatory environment, that is perhaps best addressed by subcutaneous or transdermal buprenorphine treatment as the randomized treatment.

The range of future single-patient clinical trials is almost limitless based on the rapidly increasing number of inexpensive generic medications showing efficacy in animal models. The chief constraint moving forward will be gathering experienced advice from statisticians specialized in optimization regarding identifying reliable time constants between dose adjustments or creating a central complex phenotype/endophenotype registry based on single-patient trial data [16].

## Conclusion

We found that plasma pTau181 and NfL were associated with opioid use and that pTau181 could be used to guide medication titration in clinically relevant short-time scales. Single-patient trials in patients with premature neuronal aging, where medication and dosage choices are based on individual ATN profiles, may rapidly improve treatment success.

## Acknowledgments

Authors (EH, KB) would like to thank Matt Kaeberlein and Allessandro Bitto for undergraduate training in their laboratory.

