## Supplemental Data for "Elevated plasma pTau181 in a specialty neuropsychiatric clinic is associated with opioid use and guides medication titration on a clinically relevant short-time scale"

### Supplemental Figure S1 No correlation of ATN profile biomarkers with metabolic syndrome markers

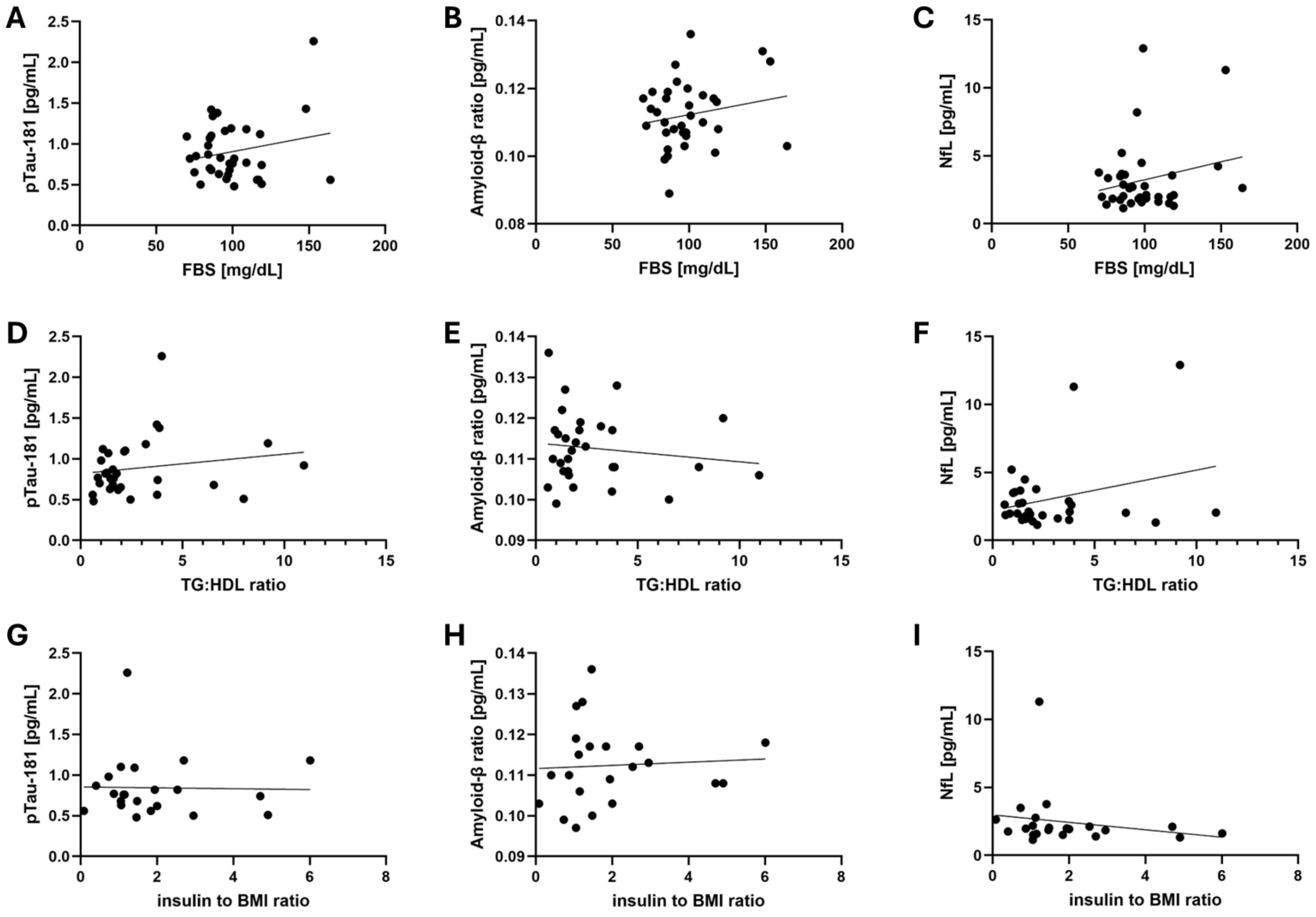

Scatter plots of the ATN profile biomarkers versus the metabolic syndrome markers. **A** pTau181 concentration versus FBS. **B** Amyloid-β ratio versus FBS. **C** NfL concentration versus FBS. **D** pTau181 concentration versus TG/HDL ratio. **E** Amyloid-β ratio versus TG/HDL ratio. **F** NfL concentration versus TG/HDL ratio. **G** pTau181 concentration versus insulin to BMI ratio. **H** Amyloid-β ratio versus insulin to BMI ratio. **I** NfL concentration versus insulin to BMI ratio.

Each black dot represents a single value in one patient. FBS was available for 35 patients, TG/HDL ratio for 30 patients, and insulin to BMI ratio for 22 patients.

ATN, amyloid-tau-neurodegeneration; BMI, body-mass index; FBS, fasting blood glucose; NfL, neurofilament light chain; pTau181, phosphorylated Tau 181; RDW, red cell distribution width; SD, standard deviation; TG:HDL, triglyceride to high-density lipoprotein.

**Supplemental Figure S2 pTau181 concentration during lithium treatment in Case #1**

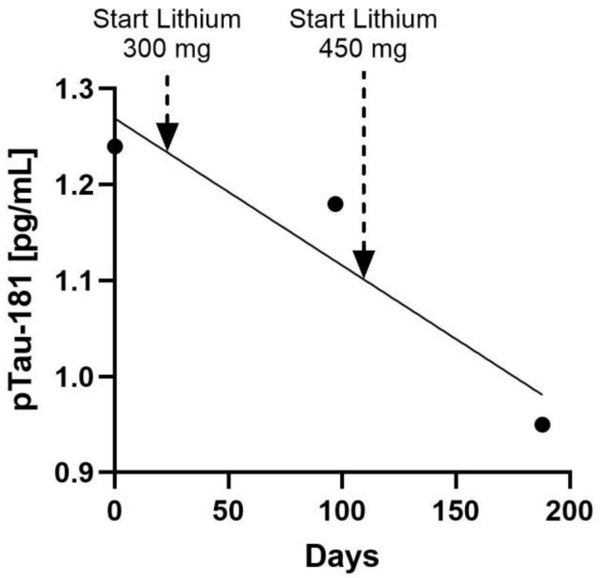


The pTau181 concentrations during lithium treatment in Case #1. The black dots indicate the concentration on the test days (day 0, day 97, day 193). Lithium 300 mg started on day 23 and lithium 450 mg started on day 107. The diagonal line indicates the simple linear regression ($r^2$=0.89).

pTau181, phosphorylated Tau 181.
